# Beta-cell adaptation to metabolic stresses requires prolactin receptor signaling

**DOI:** 10.1101/2024.01.20.575603

**Authors:** Daniel Lee, Raneet Kahlon, Darasimi Kola-Ilesanmi, Mahir Rahman, Carol Huang

## Abstract

The role of prolactin receptor (PRLR) signaling in β-cell adaptation to maternal insulin resistance of pregnancy has been well demonstrated. Using transgenic mice with an inducible β-cell-specific Prlr deletion (βPrlr^-/-^), we found that intact PRLR, as found in βPrlr^+/+^ mice, were protected from developing glucose intolerance during pregnancy, and the main mechanism responsible for this PRLR-mediated effect is the up regulation of β-cell proliferation and insulin synthesis. Interestingly, studies in male mice and humans have found a link between diminished PRLR signaling and abnormal β-cell function. We aimed to determine whether PRLR has a role in regulating β-cell function outside of pregnancy, protecting β-cell against exposure to metabolic stressors.

In this study, we found that β-cell-specific PRLR reduction resulted in impaired glucose tolerance in multiparous female mice challenged with a 12-week course of high-fat diet (HFD). Unlike in pregnancy, where PRLR signaling up regulates β-cell proliferation resulting in a greater β-cell mass, we observed no difference in β-cell mass between the wild type (βPrlr^+/+^) and mutant (βPrlr^-/-^) mice. In vitro glucose-stimulated insulin secretion using isolated islets from wild type (βPrlr^+/+^) and mutant (βPrlr^-/-^) mice showed comparable insulin response, but βPrlr^-/-^ mice showed blunted first-phase insulin release in vivo, although only when challenged with glucose orally and not intraperitoneally, suggesting an impairment of the incretin effect. In support of the observed defect in incretin action, we found a reduction in expression of both incretin hormone receptors, *Gipr* and *Glp-1r*, and several of their upstream regulators, such as *E2f1, Nkx6*.*1, Pax6, Pparγ*, and *Tcf7l2*. Islets from the mutant mice also have a lower insulin content and reduced levels of genes that regulate glucose metabolism. Together, these results suggested that PRLR signaling plays an important role in preserving β-cell function in mice exposed to metabolic stress by maintaining incretin receptor expression and insulin secretory capacity in β cells.

## Introduction

Pancreatic β cells release insulin upon nutrient intake to regulate glucose homeostasis. Diabetes ensues when insulin output fails to match insulin demand. Pregnant mothers develop insulin resistance to divert nutrients to the fetus. In order to maintain normal serum glucose, maternal pancreatic islets need to secrete more insulin to meet this increased insulin demand ^1^. Pregnancy-induced β-cell adaptation includes β-cell hyperplasia, hypertrophy, increased insulin synthesis, lowered threshold for glucose stimulated insulin secretion (GSIS), and blocking apoptosis ^2,3^. Previous work has identified prolactin receptor (PRLR) as a key regulator of β-cell proliferation, survival, and insulin release in response to the elevated insulin demand during pregnancy ^4-7^. We reported *in vivo* evidence that PRLR signaling is required for increasing β-cell mass to increase insulin synthesis capacity during pregnancy to stave off gestational diabetes (GDM) ^4,8^, mainly by up regulating β-cell replication, engaging pro-proliferative signaling molecules, namely IRS-2, pAKT, pJAK2/pSTST5, MENIN, and p21^5^. We also identified *Lrrc55* (Leucine Rich Repeat Containing 55), a putative γ-subunit of the Big Potassium channel^9^, as a novel PRLR-regulated pro-survival factor in islets ^10^. While most data on PRLR action in β cells came from studies in pregnancy, data on its role outside of pregnancy are emerging. Human studies found that postpartum prolactin level is positively associated with increased insulin sensitivity and lower prolactin levels is characterized by a lipidomic profile associated with high risk for developing type 2 diabetes (T2D) ^11^. This is consistent with epidemiological studies showing that prolactin levels in the highest quartile of normal range is associated with the lowest T2D risk, as demonstrated in both men and women, including those with a GDM history ^11-16^. Results from these studies and our previous work on the role of PRLR in β-cell adaptive responses prompted us to investigate PRLR’s role in β-cell adapation to metabolic stressors other than pregnancy, namely, exposure to a high-fat diet (HFD). Exposure to HFD, or the Western diet, is a common metabolic stress that places significant demand on β cells to secrete more insulin. In this study, we use a model of sequential exposure to metabolic stressors, i.e. 2-3 pregnancies, followed by a 12-week HFD, to mimic the human condition where women experiences multiple pregnancies, followed by consumption of a typical high-fat high calorie Western diet. Here, we report that when challenged with a 12-week course of HFD, multiparous female mice with an inducible, β-cell-specific deletion of prolactin receptor (βPrlr-/-) had impaired glucose tolerance in comparison to the wild type βPrlr+/+ mice. Unlike in pregnancy, β-cell proliferation had a minimal role in this adaptation. The main defect in the βPrlr-/- mice was a blunted in vivo first-phase insulin secretion and a decreased islet insulin content and incretin receptor expression.

## Methods

### Ethical approval

All experimental procedures were approved by the Animal Use Review Committee at the University of Calgary in accordance with standards of the Canadian Council on Animal Care.

### Mice

Generation of an inducible, β-cell-specific deletion of prolactin receptor (Prlr) (herein denoted as βPrlr-/-) was previously described ^17^. Briefly, a promoter-driven targeting cassette was obtained from EUCOMM (The European Conditional Mouse Mutagenesis Program), electroporated into mouse ES cells, injected into CD1E wild type mouse embryos to generate chimeras. After confirmation of germline transmission, mice heterozygous for floxed exon 5 of Prlr (Prlr^f+/-^) were back-crossed with C57BL/6J mice (The Jackson Laboratory) for more than 10 generations. The Prlr^+/-^ mice were crossed with Pdx1CreER™ mice (The Jackson Laboratory), and male Pdx1CreERTM: Prlr^+/-^ mice were crossed with female Prlr^+/-^ to generate the homozygous conditional knockout of βPrlr-/-(Pdx1CreER™:Prlr^-/-^), and control littermates (Pdx1CreER™: Prlr^+/+^and Prlr^+/+^). The Pdx1CreER™:Prlr^-/-^ were then crossed with the mT/mG reporter mice^18^. At age 8 weeks, 200mg/kg of tamoxifen (dissolved in corn oil) was given by oral gavage for 5 doses every other day to induce Cre recombinase activity^17^. Mice were maintained on a 12-h light, 12-h dark cycle with liberal access to food and water.

Pregnancy: Four weeks after last dose of tamoxifen, female mice were paired with wild type males. The male was removed from the cage after the female delivered 2-3 litters of pups. Two-three weeks after their pregnancies, the multiparous female mice were placed on either a HFD where 60% of calories come from fats (Research Diets D12492) or a control diet (CD) where 10% of calories come from fat (D12450K) for 12 weeks.

### Glucose homeostasis

Glucose tolerance tests -Overnight fasted mice were given glucose solution (20% D-glucose in water, 2g/kg body weight) orally (oral glucose tolerance test, or OGTT) or intraperitoneally (intraperitoneal glucose tolerance test, or IPGTT), and blood was sampled from tail vein at times 0, 10, 15, 30, 45, 60, and 120 minutes after to measure serum glucose using a glucometer (OneTouch Verio). Additional blood samples (∼30μl) were taken at times 0, 10 and 30 minutes from the saphenous vein for insulin concentration measurements by ELISA (CrystalChem, catalog number 90082). Non-fasted blood glucose was determined at 8am and 30μl of serum was taken simultaneously and stored at -80°C for measurement of insulin by ELISA.

Insulin tolerance test (ITT) - ITT was performed in early afternoon, after a 4-6 hour fast. Insulin (0.5units/kg) was administered intraperitoneally and blood glucose was sampled from tail vein at 0, 15, 30, 45, and 60 minutes after insulin injection.

### Pancreas Isolation

Pancreases were isolated using blunt dissection, cleaned of blood and fat, and weighed. Pancreases were fixed in 4% paraformaldehyde (PFA) at 4°C overnight with gentle agitation. Fixed pancreases were washed in phosphate buffered saline (PBS) and dehydrated in increasing concentrations of sucrose solutions (10, 20, and 30% sucrose in PBS). Pancreases were then preserved in optimal cutting temperature compound (Tissue-Tek O.C.T. Compound, VWR, catalog number 25608-930) and stored in -80°C for serial sectioning on a later date.

### Islet isolation

Pancreatic islets were isolated as previously described^17^. Briefly, pancreas was distended by cannulizing the bile duct to infuse 2.5ml of collagenase P (0.66mg/ml in Hank’s Balanced Salt Solution) (Roche, catalog number C7657), surgically removed and incubated at 37**°**C for 15 minutes under constant agitation. Islets were hand-picked and 20 islets were cultured overnight in RPMI 1640 with glutamine (Hyclone, catalog number SH3002701) supplemented with 10% Fetal Bovine Serum (Gibco, catalog number 12483020) and 1U/100ml Penicillin-streptomycin (Gibco, catalog number 15070063) at 37°C and 5% CO_2_ for in vitro glucose-stimulated insulin secretion assay the next day. The remaining islets were flash freeze and stored in -80°C for RNA or protein extraction on a later date.

### In vitro glucose-stimulated insulin secretion

After overnight culture, 20 islets were preincubated in Krebs-Ringer Buffer (KRB) with 2mM glucose for 30 minutes, and repeated 1-2 times. To observe basal insulin secretion, islets were transferred to microcentrifuge tubes containing 300µl of 2mM Glucose/KRB, and incubated for 1 hour at 37°C and 5% CO_2_. Following one hour of incubation, supernatant (∼280µl) was collected and stored in -80°C for insulin measurement by ELISA. 300 µl of 16mM Glucose/KRB was added to the islets and incubated in the same conditions as above, and supernatant collected after 1 hour. Finally, 300 µl of 40mM KCl/2mM Glucose/KRB was added for 1 hour and supernatant was collected. Following the 40mM KCl incubation step and removal of the supernatant, 500µl of acid-ethanol (3% HCl, 75% EtOH) was added to lyse the islets and to extract insulin. The following day, 250µl of supernatant was mixed with equal volume of 1M Tris (pH=7.5) to neutralize the HCl and stored in -80°C for insulin measurement by ELISA. Total insulin was measured as the sum of insulin in supernatant collected after 2mM Glucose, 16mM Glucose, 40mM KCl, and acid-ethanol extraction. Results are presented as the percent of insulin secreted in each media condition relative to total insulin.

### Immunostaining

The pancreas blocks were longitudinally serial sectioned to a thickness of 7 **μ**m. Every 20^th^ section was stained for insulin to identify β cells as previously described^7^. Briefly, after 1 hour of blocking with 1% goat serum/PBS at room temperature, tissues were incubated with primary antibody over night at 4**°**C (guinea pig anti-insulin at 1:4, diluted in 1% goat serum/PBS, Agilent). This was followed by 1-hour incubation with fluorophore-conjugated secondary antibodies (Cy3-anti-guinea pig, diluted in 1% goat serum/PBS at 1:300, Jackson Laboratories). Bis-benzimide H 33342 trihydrochloride (0.1**μ**g/ml, Sigma) was added to the secondary antibody for nuclear staining. Stained sections were mounted using Fluoromount-G (Southern Biotech) fluorescent mounting medium and stored at 4**°**C.

### Islet Morphometry

Consecutive images of non-overlapping, adjacent areas of the entire pancreas section were acquired using a Zeiss fluorescence microscope, and captured with a CoolSnap digital camera. Images were analyzed by ImageJ software to measure the insulin-positive area as well as the area of the entire pancreas section (identified by nuclear staining). β-cell mass was calculated by multiplying the pancreas weight by the β-cell fraction (i.e. the ratio of insulin-positive cell area to total pancreatic tissue area on the entire section). Results represent the average of 6-8 tissue sections per animal from 5-6 animals from each genotype.

### Islet RNA isolation and quantitative real-time q-PCR

Total islet RNA (200–300 islets/mouse) was extracted using the RNeasy Mini Kit (Qiagen). RNA concentration and integrity were assessed using the ND-1000 Spectrophotometer (NanoDrop). cDNA was synthesized using the Quantitect Reverse Transcription Kit (Qiagen). Primers were designed using Primer Designing Software (NCBI) (sequences available upon request). RT-qPCR reactions were carried out in triplicate with QuantiFast SYBR Green Master Mix (Qiagen) at an annealing temperature of 60ºC. Data were collected using the DNA Engine Opticon2 Continuous Fluorescence Detection System (BioRad) and software (Bio-Rad). The relative amount of RNA was determined by comparison with inorganic pyrophosphate (Ppa1) as a reference gene. It was chosen because its expression was comparable in islets from all groups.

### Statistical analysis

All statistics were performed using GraphPad Prism 4 software. Two-tailed Student’s t tests or ANOVA with Tukey’s post-tests were performed where appropriate. Comparisons were made between β-cell-specific Prlr-deletion mutant mice (βPrlr-/-) and the wild type littermate (βPrlr+/+), as stated in the Figure Legend.

## Results

### Intact PRLR is required for maintenance of normal glucose homeostasis in multiparous female mice exposed to a high-fat diet

Prolactin receptor (PRLR) has been shown to be important for regulating glucose homeostasis during pregnancy in both whole body ^4^ and β-cell specific Prlr-knockout mice ^17,19^, mainly by up regulating β-cell proliferation and increase insulin synthesis and release to compensate for the insulin resistance of pregnancy. To determine whether PRLR also plays a role in regulating glucose homeostasis and β-cell function outside of pregnancy, we exposed multiparous female mice with an inducible, β-cell-specific conditional knockout of prolactin receptor (βPrlr-/-) and their wild type (βPrlr+/+) littermates to 12 weeks of HFD (Figure 1). We found that after 6 weeks of HFD, the βPrlr-/- mice had higher glucose excursion during an oral glucose tolerance test (OGTT) than their wild type littermates, a difference that persisted until the end of the 12-week HFD period (Figure 2a-c). There was no difference in fasting blood glucose between βPrlr+/+ and βPrlr-/- mice after 6 or 12 weeks of HFD (6 weeks -βPrlr+/+: 9.17±0.68 mM, βPrlr-/-: 7.85±0.64 mM, p=0.10; 12 weeks – WT:7.65±0.61 mM, βPrlr-/-: 8.61±0.67mM, p=0.46, n=14-20). We also observed no significant difference in non-fasting blood glucose between the two groups throughout the 12-weeks of HFD (Figure 2d). The higher glucose excursion observed in the βPrlr-/- mice was not due to a difference in insulin sensitivity, as we observed comparable drop in serum glucose during an insulin tolerance test (ITT) between the two groups (Figure 2e).

**Figure 1.**
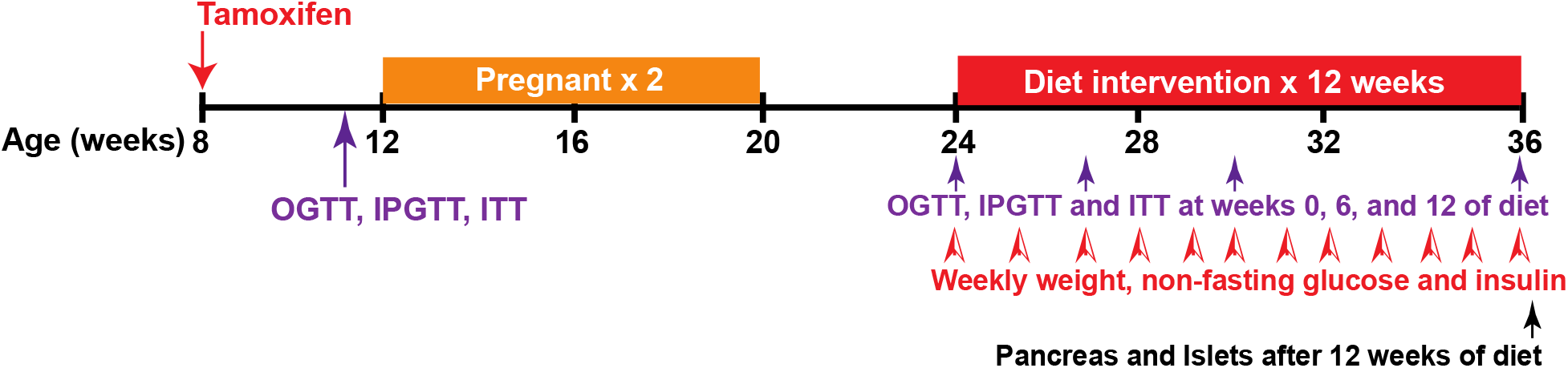
Experimental Design. Both wild type and βPrlr-/-female mice were given tamoxifen at age 8 weeks, and 4 weeks later, set up for 2-3 pregnancies, then placed on a 12-week course of HFD or control diet (CD). Glucose homeostasis was measured by oral glucose tolerance test (OGTT), intraperitoneal tolerance test (IPGTT), and insulin tolerance test (ITT) at times indicated.

**Figure 2.**
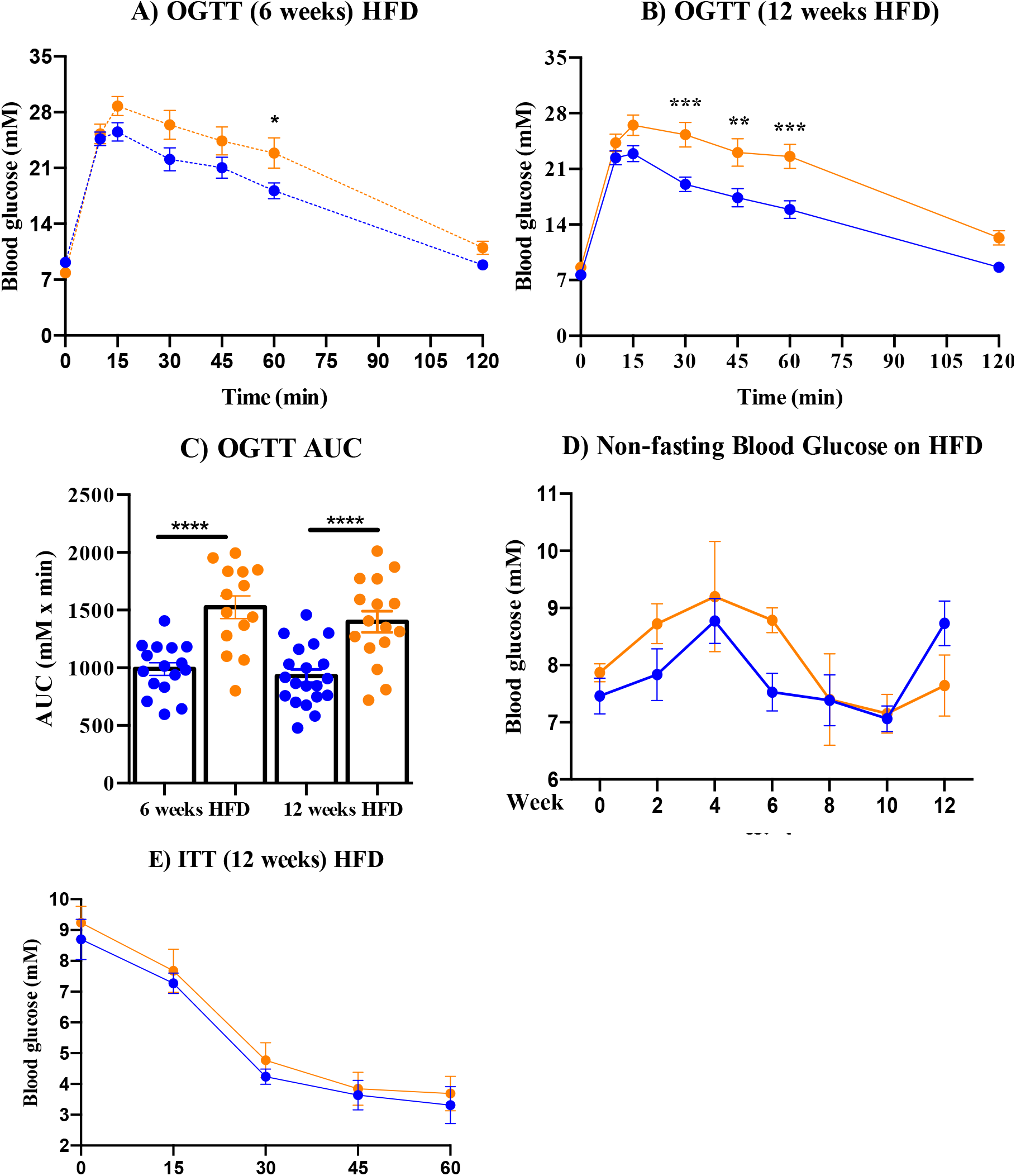
Multiparous female βPrlr-/- mice had impaired oral glucose tolerance. Glucose levels throughout an OGTT at week 6 (A) and week 12 (B) of the 12-week HFD course. Glucose excursions, measured as integrated area under the curve (AUC) across all time points throughout the 120-minutes of OGTT, are presented (C). Non-fasting blood glucose (D) were taken at 8am throughout the 12-week HFD period. (E) Insulin tolerance test was performed after an 4-6 hour fast. Results are expressed as mean + SEM. One-way ANOVA with Tukey’s post hoc test was performed to compared between βPrlr+/+ and βPrlr-/-: “*”= *p<0*.*05*,“**”=*p<0*.*005*,“***”=*p<0*.*0005*,“****”= *p<0*.*00005*. For A-D, n=14-20 mice for each group; for E: n=3-18 mice/group. Blue = βPrlr+/+ mice, Orange = βPrlr-/- mice.

### βPrlr-/- mice secreted less insulin during an OGTT

To understand the cause of the higher glucose excursion in the βPrlr-/- mice, we measured in vivo insulin secretion during an OGTT. We found no difference in serum insulin levels at time 0 or 30 minutes of the OGTT, but the βPrlr-/- mice secreted significantly less insulin at the 10-minute time point (Figure 3a). Insulinogenic index, calculated as the change in insulin concentration over the change in glucose concentration, was also blunted at the 10-minute time point in the βPrlr-/- mice in comparison to the βPrlr+/+ mice. Interestingly, in vitro GSIS using isolated islets showed no difference in insulin secretory responses (Figure 3b). This suggests that islets have no intrinsic defect in glucose-stimulated insulin secretion that can account for the impaired first-phase insulin secretory response observed in vivo in the βPrlr-/- mice.

**Figure 3.**
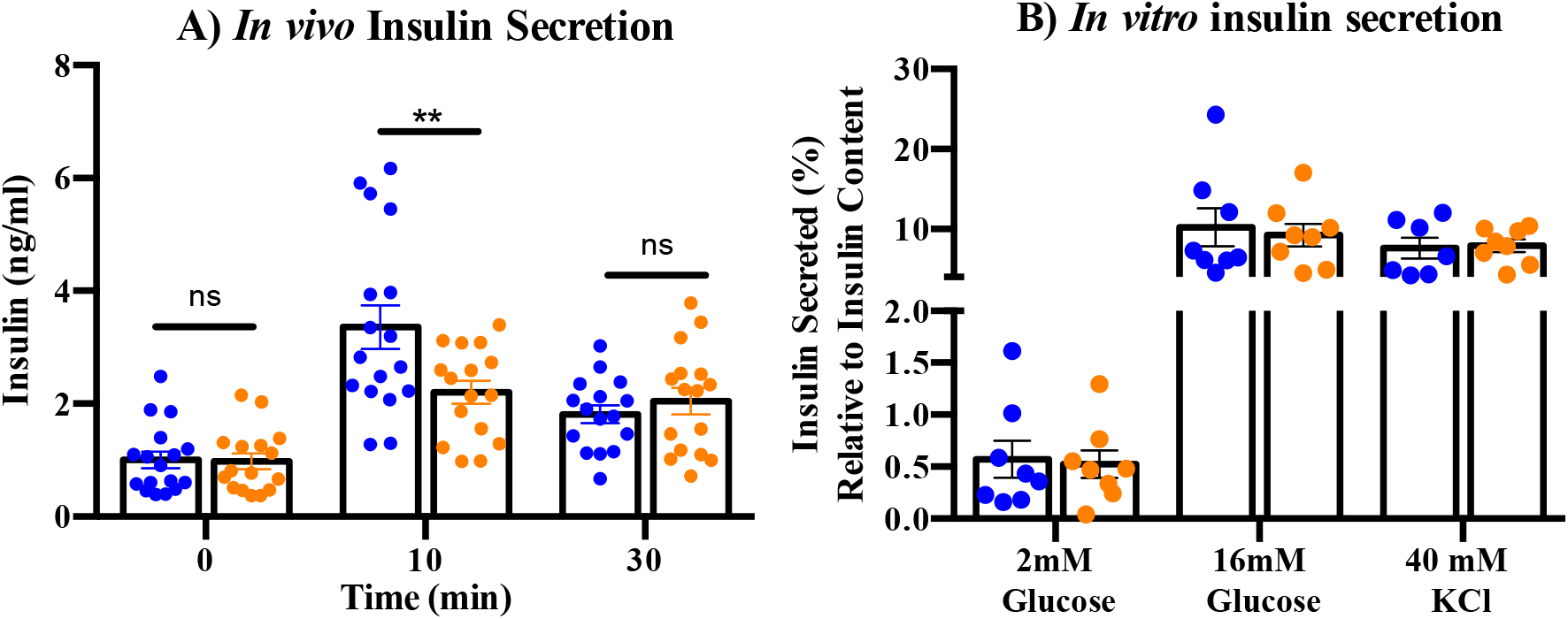
Insulin secretion after 12 weeks of HFD. (A) Plasma insulin concentrations during oral glucose tolerance tests (2g/kg) in βPrlr+/+ and βPrlr-/- mice after 12 weeks of HFD. Blood was collected at 0, 10, and 30 minutes and measured by ELISA. Results are expressed as mean + SEM (n=16-17 mice/group). (B) *In vitro* glucose-stimulated insulin secretion from isolated islets cultured in 2mM and then 16mM glucose, followed by 40 mM KCl. Results were normalized to total insulin content. Results are expressed as mean + SEM; (n = 8 mice/group). Statistical analysis was performed by one-way ANOVA with Tukey’s post hoc test where “*\*\*”=p<0*.*005*βPrlr+/+ versus βPrlr-/- mice. “ns” = not significant.

### There is little difference in β-cell proliferation, apoptosis or mass between the βPrlr-/-and βPrlr+/+ mice

During pregnancy, it has been well established that intact Prlr signaling is required for β-cell adaptation to the insulin resistance of pregnancy by up regulating β-cell proliferation, increasing β-cell mass and boosting insulin secretion^3,4,19^. To examine whether the same mechnism is responsible for the impaired glucose tolerance observed in βPrlr-/- mice after a 12-week HFD, we measured β-cell proliferation and apoptosis rates by staining β cells for ki67 and cleaved caspase 3, respectively. In contrast to our previous observation in pregnancy, where Prlr deletion significantly impacted β-cell proliferative capacity, we observed a small difference in β-cell proliferation and apoptosis rates with no significant difference in β-cell mass between βPrlr+/+ and the βPrlr-/- mice (Figure 4). Therefore, the difference in glucose tolerance cannot be explained by a difference in β-cell mass, as was the case in pregnancy.

**Figure 4.**
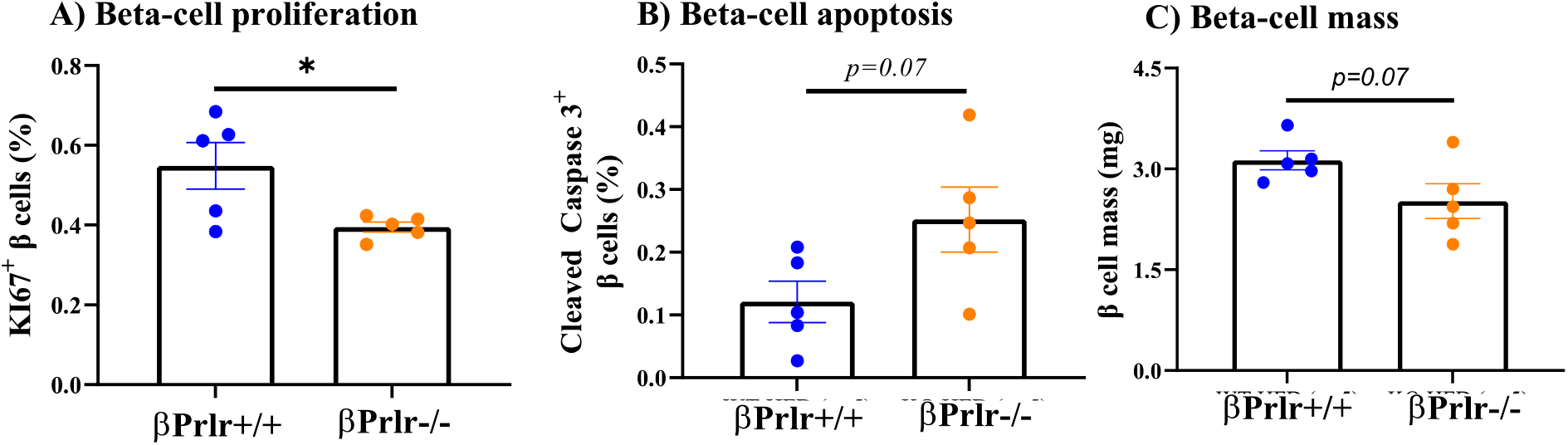
Beta-cell mass after 12 weeks of HFD. (A) Proliferating β cells were identified by ki67 and insulin double positivity; at least 6000 β cells/mouse were counted. (B) Apoptotic β cells was identified by cleaved caspase 3 and insulin double positivity; at least 3000 β cells/mouse were counted. (C) β-cell mass determined from at least three sections and 6000 β cells per mouse. Each data point represents a mouse (n = 5 mice/group). Statistical analysis was performed using student’s t-test where **“*”=** *p < 0*.05 between βPrlr+/+ and βPrlr-/- mice.

### βPrlr-/- mice had a lower islet insulin content and reduced incretin receptor expression

To identify potential mechanism contributing to the blunted in vivo GSIS observed in βPrlr-/- mice, we measured insulin content of isolated islets. We found that in comparison to islets from βPrlr+/+ mice, islets from βPrlr-/- mice had significantly lower insulin content (βPrlr+/+: 4.4±0.8μg, βPrlr-/-: 1.8±0.2μg of insulin, normalized to DNA content, p=0.03)(Figure 5a). This was accompanied by a reduction in expression of insulin 1 (*Ins1*) and insulin 2 (*Ins2*) genes (Figure 5b). We measured expression of *MafA, NeuroD1, Pdx1*, and *Nkx6*.*1*, genes that have been shown to regulate insulin gene transcription, and found a reduction in expression of all 4 genes, although only *Nkx6*.*1* reached statistical significance (Figure 5c). We also measured expression of genes that regulate GSIS^20^, namely glucose transport 2 (*Glut2*)^21^, glucokinase (*Gck*)^22^, and pyruvate carboxylate (*PC*)^23^, and found that their expression were also blunted in the βPrlr-/- mice (Figure 5d). Next, we determined the expression of incretin receptors, since incretins are potent stimulant of in vivo GSIS. Here, we found that expression of both glucagon-like peptide 1 receptor (*Glp-1r*) and glucose-dependent insulinotropic polypeptide receptor (*Gipr*) are lower in the βPrlr-/- mice (Figure 5e), and genes that regulate *Glp-1r* and *Gipr* expression, namely *E2f1*^24^, *Nkx6*.*1*^25^, *Pax6*^26^, *Pparγ*^27^, and *Tcf7l2*^28,29^ were all down regulated in the βPrlr-/- mice (Figure 5f), providing a potential mechanism for the blunted in vivo first-phase insulin secretion observed in the βPrlr-/- mice. This suggests a blunted incretin action, supported by our observation that glucose excursion was comparable during an IPGTT between βPrlr+/+ and βPrlr-/- mice (data not shown) while glucose excursion was higher in βPrlr-/- mice during an OGTT, as OGTT but not IPGTT induces a robust glucose-stimulated incretin release and incretin-augmented insulin release. Moreover, while we observed a blunted insulin response at time 10-minute of OGTT in βPrlr-/- mice (Figure 3a), there was no difference in plasma insulin levels at any time point throughout an IPGTT (Figure 5g). Expression of a downstream effector of incretin hormone receptors, Epac1, was decreased in the βPrlr-/- mice (Figure 5h) and\ incretin-mediated nuclear translocation^30^ of Pdx1 was reduced (Figure 5i).

**Figure 5.**
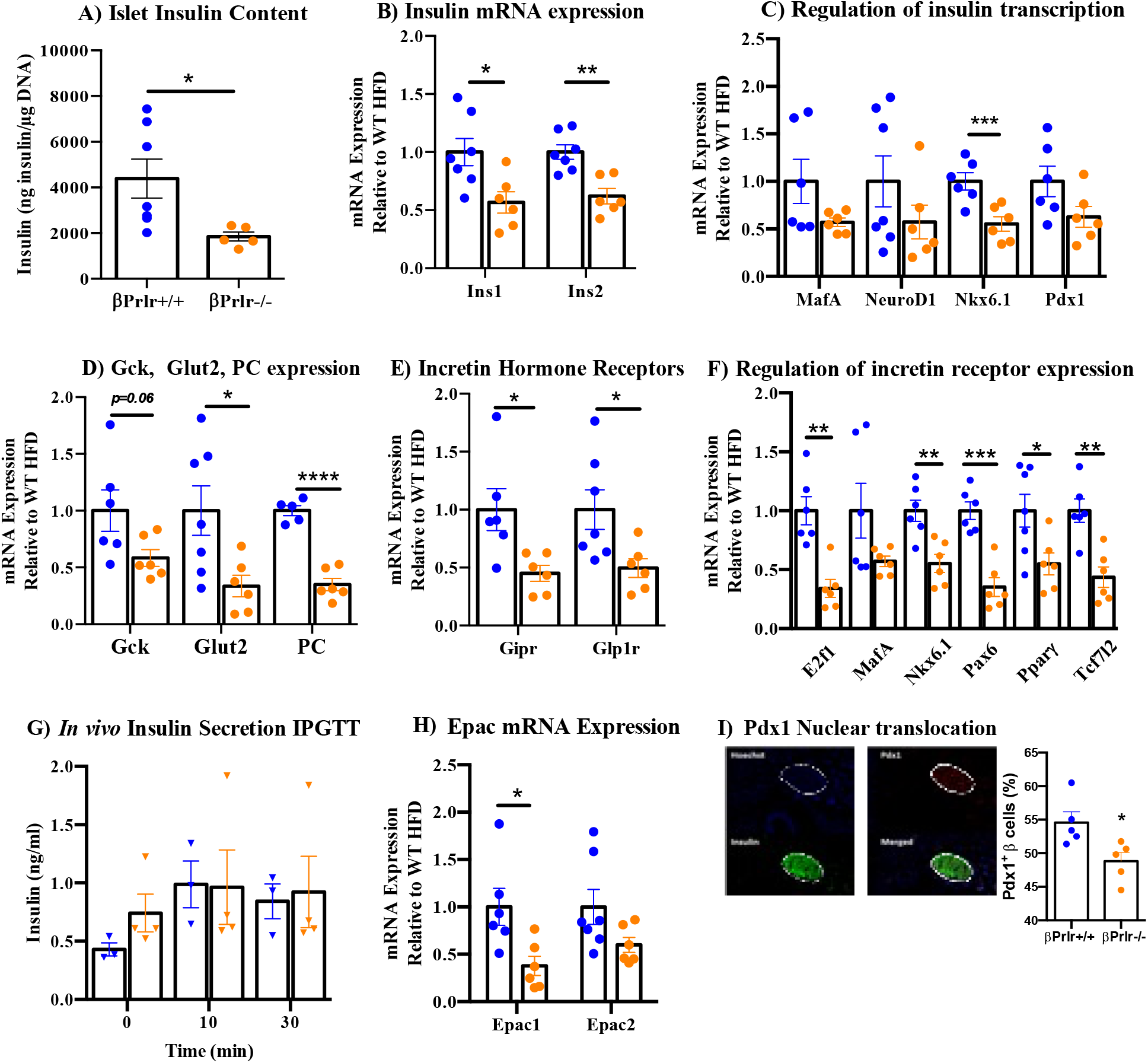
Insulin content and incretin hormone receptor expression are lower in islets of βPrlr-/- mice. **(A)** Insulin content from pancreatic islets was normalized to DNA (in μg) and expressed as mean + SEM; n = 5-7 mice/group. mRNA expression of the (B) insulin genes (*Ins1* and *Ins2*), (C) transcription factors that regulates insulin gene transcription (*MafA, NeuroD1, Nks6*.*1, Pdx1*), (D) genes that regulates glucose entry and metabolism in β cells (glucokinase (*Gck*), glucose transporter 2 (*Glut2*), pyruvate carboxylase (*PC*)), (E) incretin hormone receptors (glucagon-like peptide-1 receptor (*Glp-1r*), glucose-dependent insulinotropic polypeptide receptor (*Gipr*)), and, (F) transcription factors that regulates incretin hormone receptor expression (*E2f1, MafA, Nkx6*.*1, Pax6, Pparγ, Tcf7l2*) in islets were determined by RT-qPCR, normalized to *Ppa1* (housekeeping gene), and expressed relative to the levels found in βPrlr+/+/HFD mice. Each data point represents one mouse (n=6-7 mice/group), averaged from 3 independent experiments. Results are presented as means + SEM Statistical analysis was done using an unpaired student’s t-test between groups where “*”= *p<0*.*05, “*\*\*”= *p<0*.*005*. (G) Plasma insulin concentrations during IPGTT (2g glucose/kg body weight) in βPrlr+/+ and βPrlr-/- mice after 12 weeks of HFD. Blood was collected at 0, 10, and 30 minutes and measured by ELISA. Results are expressed as means + SEM (n=16-17 mice/group). ANOVA and Tukey’s post hoc test were performed.“*”=*p<0*.*05*,“*\*\*”= p<0*.*005* OGTT vs. IPGTT, and ns=not significant.

## Discussion

Physiologic states such as pregnancy, obesity, and aging are characterized by insulin resistance, and pancreatic beta cells adapt by increasing insulin synthesis and secretory capacity to maintain normal glucose homeostasis^31^. Previously, studies in mice with global^4^ or β-cell-specific prolactin receptor (Prlr) deletion ^17,19,32^ demonstrated that PRLR is required for β-cell adaptation to insulin resistance of pregnancy, mainly by up regulating β-cell proliferation. This adaptation involves activity of many pathways^2^, such as MafB^19^, Jak2/Stat5^5,33^, IRS-2/PI3K/Akt^5,34^, Hnf4-a^35^, menin/p27/p18^5,36^, Tph1^37^, Foxd3^38^, and FoxM1^39^. Little is known about whether PRLR has a role in regulating β-cell function outside of the context of pregnancy. Here, we report that in female multiparous mice exposed to a HFD, PRLR is important in regulating β-cell function and maintaining glucose homeostasis. In our model, female βPrlr+/+ and βPrlr-/- mice were placed on a HFD or CD for 12 weeks after they have given birth to 2-3 litters. This was to mimic the human experience of multiple pregnancies and consumption of a typical high-fat Western diet. We deliberately did not start the HFD before pregnancies since the effect of HFD on islet function during pregnancy has been well studied, and our model of HFD only after pregnancies mimics women who maintain a healthier diet during pregnancy followed by consumption of a typical Western diet after pregnancy.

This study utilized a transgenic mouse model with a β-cell-specific homozygous deletion of Prlr (βPrlr-/-) to study its role in regulation of β-cell function^17^. Female βPrlr-/- mice have impaired glucose tolerance on day 15 of pregnancy^17^; however, we did not detect a difference in glucose tolerance between virgin βPrlr+/+ and βPrlr-/- mice or after their second pregnancy, before we placed them on HFD (data not shown). This suggests that while PRLR action is required for β-cell adaptation during pregnancy, the lack of PRLR action during pregnancy do not cause permanent functional defect in β cells, as indicated by normal glucose tolerance and insulin secretion postpartum. This is consistent with our previous observation that while expression of ER stress markers are up regulated during pregnancy^10^, it did not translate into increased β-cell apoptosis during pregnancy in mice with global Prlr deletion^4^.

Following two pregnancies, mice were placed on either CD or HFD for the duration of 12 weeks. Diet induced obesity (DIO) model is most often used to mimic Western diet, which is characterized by high fat content^40-43^. CD did not affect glucose homeostasis in the βPrlr+/+ mice, but the βPrlr-/- mice had higher blood glucose levels at the 45-minute time point of OGTT, although we did not observe a significant difference in glucose excursion when measured integrated area-under-curve (AUC) throughout the 120-minutes OGTT (data not shown).

The physiological effects of HFD in rodents has been studied extensively, and it includes increasing body weight, worsened glucose tolerance, and becoming more insulin resistant ^41,44,45^. The βPrlr+/+ and βPrlr-/- mice had comparable changes in weight over the course of 12 weeks of CD or HFD. Glucose tolerance worsened in both βPrlr+/+ and βPrlr-/- mice after 6 and 12 weeks of HFD, but in comparison, the βPrlr-/- mice were more glucose intolerant than the βPrlr+/+ mice after 6 weeks of HFD, a difference that persisted until end of the 12-week HFD period (Figure 2a-c). Both βPrlr+/+ and βPrlr-/- mice became progressively more insulin resistant during HFD, as evident from their elevated blood glucose at the 15-minute time point of ITT (data not shown). These results suggest that while both βPrlr+/+ and βPrlr-/- mice experienced worsened glucose tolerance and insulin sensitivity after exposure to HFD, βPrlr-/- mice are more glucose intolerant with similar levels of insulin sensitivity in comparison to the βPrlr+/+ mice, suggests a reduction in insulin secretion in the βPrlr-/- mice.

Exposure to high fat has been shown to negatively impact insulin secretion through several mechanisms. In vitro studies found that saturated fat activates pERK1/2, stimulates *ATF6* cleavage, down regulates *MafA* and *Pdx1*, and caused nuclear exclusion of Pdx1, impairing β-cell function and inhibiting insulin gene transcription ^46-49^. Quantification of insulin content from isolate islets revealed that islets from βPrlr-/- mice given a 12-week HFD had a lower insulin content in comparison to βPrlr+/+ mice (Figure 5a). In addition, they had a reduction in mRNA expression of both *Ins1* and *Ins2* genes (Figure 5b). Of the key transcription factors that are known to directly regulate insulin gene transcription, namely *MafA, NeuroD1, Nkx6*.*1*, and *Pdx1*, we observed a reduction in expression of all 4 genes although only *Nkx6*.*1* reach statistical significance (Figure 5c). We then investigate genes involved in β-cell function that enhance GSIS^20-22^, i.e. *Glut2, Gck*, and *PC*. First, Glut2 is the only glucose transporter expressed in β cells with a high Michaelis constant (*K*_m_) and transport ability, allows quick glucose entry into the β-cell. Once glucose enters, it is phosphorylated by the rate-limiting enzyme Gck, which plays the pivotal role in GSIS. Finally, pyruvate can enter the mitochondrial tricarboxylic acid (TCA) cycle through two pathways controlled by either pyruvate carboxylase (PC) or pyruvate dehydrogenase (PDH). In β cells, evidence of PC rather than PDH, controlling pyruvate entry into the TCA cycle has been shown^50^. Here, we found a reduction in expression of *Glut2* and *PC* (Figure 5d). Hence, islets from βPrlr-/- mice appears to have widespread albeit modest reduction in genes that regulate insulin gene transcription and GSIS. In search of other potential cause of impaired insulin secretion in βPrlr-/- mice, we examined expression of pro-survival and pro-apoptotic genes. In vivo studies showed that upon HFD exposure, pro-survival gene expression increases within the first 2 weeks, followed by activation of UPR and pro-apoptosis genes^43,51^. This sequential activation of prosurvival genes and UPR pathway is also observed in β cells during pregnancy^10^. Prolactin has been shown to protect β cells from glucolipotoxicity^52^ and in our multiparous female βPrlr-/- mice given a HFD, examination of the UPR pathway revealed no significant difference in the expression of *Bax:Bcl-2* ratio, spliced *Xbp1*: unspliced *Xbp1*, and *Chop*, but the expression of *Bip* and *Ire1α* was decreased. We observed a significant reduction in the pro-survival gene, Lrrc55^10^, in the βPrlr-/- mice in comparison to the βPrlr+/+ mice. These differences were accompanied by a slightly higher β-cell apoptosis rate in the βPrlr-/- mice. We also found a small reduction in β-cell proliferation rate in the βPrlr-/- mice. However, this small changes in β-cell apoptosis and proliferation rates did not result in a statistically significant difference in β-cell mass after 12 weeks of HFD (Figure 4c). These findings suggest that unlike pregnancy, the blunted insulin secretory response in the βPrlr-/- mice was due to a defect in β-cell proliferation resulting in a smaller β-cell mass, a different compensatory mechanism is responsible for the difference in insulin secretion in multiparous mice exposed to HFD. This is in line with transcriptomic analyses where prolactin-induced genes in islets during pregnancy and HFD showed very little overlap, suggesting that these metabolic stressors activate different mechanisms of compensation^40^.

With this difference in mind, we measured insulin secretion *in vivo* and *in vitro*. We found that *in vivo* insulin secretion and the insulinogenic index was decreased at the 10-minute time point of an OGTT, suggesting a reduction in first-phase insulin secretion (Figure 3a). Curiously, this difference was not observed during an IPGTT. Moreover, we found no difference in *in vitro* insulin secretion in response to glucose-dependent (16 mM Glucose) or glucose-independent (40mM KCl) stimuli (Figure 3b).^22^ In vivo insulin secretion results from integration of nutrient signals, such as glucose, free fatty acids, amino acids, as well as hormones and neuronal signals, while in vitro GSIS are independent from these systemic effects. Our results suggest that there is no intrinsic defect in βPrlr-/-islets’ ability to secrete insulin in response to glucose but rather, the impaired first-phase insulin secretion observed *in vivo* is secondary to an in vivo factor. One such potential in vivo factor is the incretin hormones^53^. The incretin hormone receptors GIPR and GLP-1R were of interest as the incretin effect accounts for up to 80% of insulin secretion in response to meal ingestion^53^. We observed a significant reduction in *Gipr* and *Glp-1r* gene expression in islets of βPrlr-/- mice (Figure 5e), as well a reduction in *Epac1*, a downstream mediator of *Gipr* and *Glp-1r* action^54^. Epac proteins increase intracellular Ca^2+^ to promote insulin granule exocytosis and decreased expression directly impairs the incretin effect^55,56^.

Activation of both incretin hormone receptors results in the nuclear translocation of Pdx1 and we observed a reduction in number of nuclear Pdx1^+^ β cells in the βPrlr-/- mice (Figure 5i). We also measured the expression of transcription factors that regulate *Gipr* and/or *Glp-1r* expression, namely *E2f1, MafA, Nkx6*.*1, Pax6, Pparγ*, and *Tcf7l2*, and found a significant reduction in the expression of all except *MafA* when comparing βPrlr-/-to βPrlr+/+ mice (Figure 5f). It is interesting to note that all of these transcription factors have been associated with β-cell adaptation during HFD^43,57-59^, but only *E2f1* and *Pparγ* are known to be downstream of PRLR signaling^60,61^.

Hence, our results showed that during pregnancy, PRLR signaling up regulates β-cell proliferation and insulin synthesis in adaptation to the insulin resistance of pregnancy^4,17,19,40^. In the absence of additional metabolic challenges, a reduction in PRLR signaling in β cells have no impact on β-cell function and glucose homeostasis. However, when β cells are Prlr deficient, as in βPrlr-/- mice, the stress of repeated pregnancies compounded by exposure to HFD resulted in a reduction in insulin synthesis, expression of the genes that regulate GSIS, and expression of incretin receptors *Glp-1r* and *Gipr* that together, manifested as a reduction in *in vivo* insulin secretion and impaired glucose tolerance. The link between PRLR and incretin receptor expression is a novel finding and future experiments will delineate the mechanisms involved.

